# Visualization of giant virus particles using BONCAT labeling and STED microscopy

**DOI:** 10.1101/2020.07.14.202192

**Authors:** Mónica Berjón-Otero, Sarah Duponchel, Thomas Hackl, Matthias Fischer

**Affiliations:** Max Planck Institute for Medical Research, Heidelberg, Germany

**Keywords:** Cafeteria roenbergensis virus, CroV, Giant virus, *Cafeteria burkhardae*, bioorthogonal noncanonical amino acid tagging, BONCAT, L-azidohomoalanine, AHA

## Abstract

Giant DNA viruses of the phylum *Nucleocytoviricota* are being increasingly recognized as important regulators of natural protist populations. However, our knowledge of their infection cycles is still very limited due to a lack of cultured virus-host systems and molecular tools to study them. Here, we apply bioorthogonal noncanonical amino acid tagging (BONCAT) to pulse label the marine heterotrophic flagellate *Cafeteria burkhardae* during infection with the lytic giant virus CroV. In absence of CroV, we report efficient incorporation of the L-methionine analog L-azidohomoalanine (AHA) into newly synthesized proteins of the methionine prototrophic *C. burkhardae*. During CroV infection, AHA was predominantly found in viral proteins, and single CroV virions were imaged with stimulated emission depletion (STED) super-resolution microscopy. CroV particles incorporated AHA with 95-100% efficiency while retaining their infectivity, which makes BONCAT/STED a powerful tool to study viral replication cycles in this ecologically relevant marine bacterivore.

**Significance:** Giant DNA viruses are the dominant class of protist-infecting viruses, yet the vast majority of described giant virus-protist systems remain uncultured. One of the better studied cultured systems is composed of the stramenopile *Cafeteria burkhardae* (previously *C. roenbergensis*), the giant Cafeteria roenbergensis virus (CroV) and the virophage mavirus. *C. burkhardae* is a widespread marine phagotrophic protist that plays an important role in regulating bacterial populations. In addition to being grazed upon by larger zooplankton, *C. burkhardae* populations are controlled by the lytic giant virus CroV. In turn, CroV is parasitized by the virophage mavirus that increases host population survival in the presence of CroV and forms a mutualistic symbiosis with its host. Despite being of fundamental ecological and evolutionary interest, this tripartite host-virus-virophage system suffers from a lack of molecular tools. Here, we show that CroV particles can be fluorescently labeled and imaged by super-resolution microscopy. To achieve this we established robust procedures for analyzing protist and viral populations and implemented the use of bioorthogonal noncanonical amino acid tagging (BONCAT) in a marine unicellular flagellate.

## Introduction

*Cafeteria roenbergensis* is a marine unicellular heterotrophic biflagellated protist with a cell diameter of 2 μm to 6 μm (Fenchel and Patterson, 1988; O’Kelly and Patterson, 1996). It belongs to the order Bicosoecida (Stramenopiles) and is widely distributed in marine environments such as surface waters, deep sea sediments, and hydrothermal vents (Atkins *et al.*, 2000; Scheckenbach *et al.*, 2005). As bacterivores, *Cafeteria* flagellates play an important role in carbon transfer and nutrient recycling (Boenigk and Arndt, 2002; Pernthaler, 2005). Recently, many of the circulating *C. roenbergensis* strains have been reclassified as *C. burkhardae* based on 18S rDNA analysis (Schoenle *et al.*, 2020), including the strains used in this study. Studies on *Cafeteria* sp. have been mainly focused on bacterial grazing and its ecological impact (Otto *et al.*, 1997; Boenigk and Arndt, 2000; Zubkov and Sleigh, 2000; De Corte *et al.*, 2019), whereas the molecular biology of this Stramenopile remains largely unknown. We recently published the first genome assemblies for *C. burkhardae* which showed that the 35 Mbp genome is mostly diploid (Hackl *et al.*, 2020). In the wild, *Cafeteria* populations are modulated by their interactions with viruses (Massana *et al.*, 2007). Two of these, Cafeteria roenbergensis virus (CroV) and mavirus, have been isolated from *Cafeteria* cultures in the past, revealing a fascinating tripartite host-virus-virophage system (Garza and Suttle, 1995; Fischer *et al.*, 2010; Fischer and Suttle, 2011).

CroV is a lytic giant virus of the family *Mimiviridae* (Fischer *et al.*, 2010). CroV particles are 300 nm in diameter and composed of an outer protein shell with icosahedral symmetry, an internal lipid membrane, and a proteinaceous core (Xiao *et al.*, 2017). The ≈700 kilobase pair double-stranded DNA genome encodes more than 500 predicted proteins (Fischer *et al.*, 2010, 2014) including DNA replication and transcription machineries, which enables CroV to replicate in the host cytoplasm, in the so-called virion factory, similar to other mimiviruses and also poxviruses (Schramm and Krijnse-Locker, 2005; Suzan-Monti *et al.*, 2007; Mutsafi*et al.*, 2010). Although CroV encodes some translation components, it lacks ribosomal proteins and therefore depends on host translation. Newly synthesized CroV particles are released as early as 5-6 hours post infection (hpi), while the majority of extracellular virions appear during host cell lysis at 15-22 hpi (Taylor *et al.*, 2018).

The second virus in the tripartite *Cafeteria* system is the virophage mavirus of the family *Lavidaviridae*, an obligate parasite that depends on the presence of the CroV for its propagation (Fischer and Suttle, 2011; Krupovic *et al.*, 2016). During a co-infection of *C. burkhardae*, mavirus severely reduces CroV particle production thereby protecting the still CroV-uninfected part of the host population (Fischer and Hackl, 2016). In addition, mavirus enters *C. burkhardae* cells independently of CroV and integrates its genome into the host genome, where virophage gene expression does not occur until the cell, or one of its descendants carrying the endogenous mavirus, is infected with CroV. Once reactivated by CroV, mavirus replicates and CroV and mavirus particles are released by cell lysis (Fischer and Hackl, 2016; Berjón-Otero *et al.*, 2019; Duponchel and Fischer, 2019). Mavirus may thus provide an anti-viral defense for *C. burkhardae* populations against the spread of CroV.

Currently, the lack of molecular tools hampers a deeper mechanistic understanding of this tripartite system. We, therefore, explored bioorthogonal noncanonical amino acid tagging (BONCAT) as a fast and easy technique for protein labeling during viral infection. BONCAT is based on the incorporation of a noncanonical amino acid into nascent polypeptide chains during protein translation (Kiick *et al.*, 2002; Dieterich *et al.*, 2006). One of these noncanonical amino acids is L-azidohomoalanine (AHA), a nontoxic and water-soluble analog of methionine containing an azido moiety. Thus, when AHA is present in the culture medium, cellular uptake and loading by methionyl-tRNA synthetase lead to AHA incorporation into newly synthesized proteins. AHA-containing proteins can then be purified via an alkyne-bearing biotinylated tag, or coupled to a fluorophore via Cu(I)-catalyzed azide alkyne cycloaddition (CuAAC), also known as “click chemistry” or Click-iT^®^ (Rostovtsev *et al.*, 2002; Tornøe *et al.*, 2002; Zhang *et al.*, 2017). To maximize the incorporation rates of the noncanonical amino acid, this technique is often applied in methionine auxotrophic organisms or cell culture systems such as mammalian cells (Dieterich *et al.*, 2007; Zhang *et al.*, 2017). However, in recent years, efficient AHA incorporation has been demonstrated in natural samples (Hatzenpichler *et al.*, 2014; Leizeaga *et al.*, 2017). BONCAT can also be used to label viral proteins when AHA or another noncanonical amino acid is added during viral infection (Banerjee *et al.*, 2011; Huang *et al.*, 2017; Wang *et al.*, 2020).

Here we show successful BONCAT labeling of newly synthesized proteins in *C. burkhardae*, a eukaryotic organism containing essential genes for methionine biosynthesis. When this protist is infected with CroV, AHA is incorporated in viral proteins allowing the visualization of giant virus particles with high-resolution microscopy. AHA incorporation in CroV particles is highly efficient (95-100%) and AHA does not impair the infectivity or virion yield of CroV. Our results prove that antibody-free metabolic labeling combined with super-resolution microscopy is an effective method to visualize giant viruses by fluorescence microscopy. The protocols described here provide the basis for future microscopy studies of *C. burkhardae* and its viruses.

## Results and discussion

### Essential methionine biosynthesis genes are present and transcribed in *C. burkhardae*

Because high endogenous methionine levels can impair efficient BONCAT labeling with methionine analogs such as AHA, methionine auxotrophic organisms are preferably used for this technique (Wisse *et al.*, 2017; Zhang *et al.*, 2017). Therefore we first examined the methionine biosynthesis coding potential in *C. burkhardae* using the recently published genome assembly of *C. burkhardae* strain RCC970-E3 (Hackl *et al.*, 2020). We queried all predicted protein sequences encoded by *C. burkhardae* against the methionine pathway proteins as described in the diatom *Thalassiosira pseudonana* (Bromke and Hesse, 2015). BLAST analysis revealed homologs for all genes in question (Fig. 1A and Table S1). Interestingly, three of these enzymes (aspartate kinase, GenBank acc. numbers KAA0156511 and KAA0162106; homoserine acetyltransferase, KAA0152884 and KAA0165906; methionine synthase, KAA0150236 and KAA0158901) were represented by two separate genes each in the *C. burkhardae* genome. The presence of two non-homologous aspartate kinases was also reported in *T. pseudonana* (Bromke and Hesse, 2015), also two methionine synthases were found in other eukaryotes such as *Phaeodactylum tricornutum* (Helliwell *et al.*, 2011), however non-homologous homoserine acetyltransferases, to the best of our knowledge, have so far only been described in bacteria (Ferla and Patrick, 2014).

**Figure 1.**
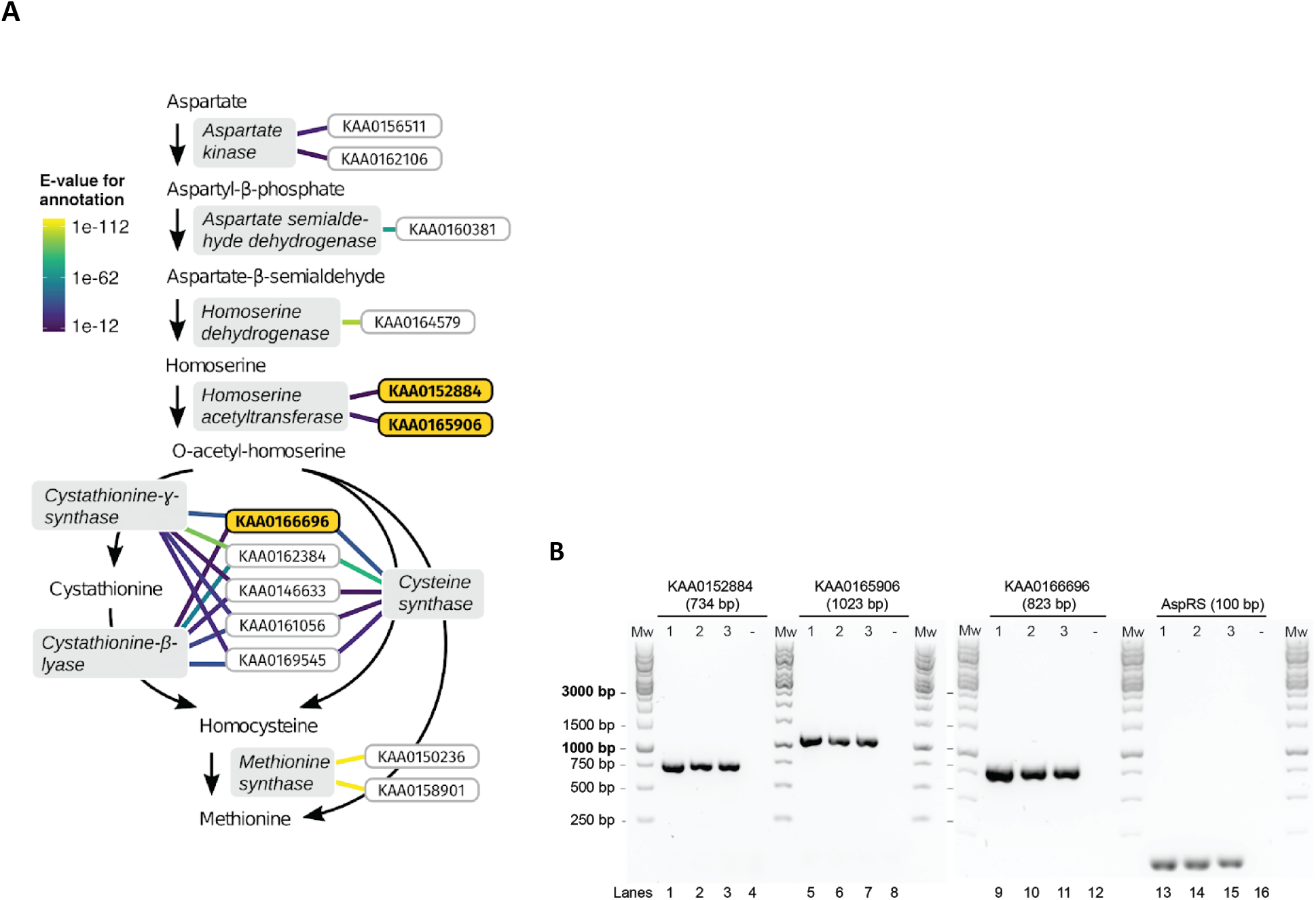
The inferred methionine biosynthesis pathway in *C. burkhardae*. (A) Gene products of *C. burkhardae* predicted to function in methionine biosynthesis are listed with their GenBank accession numbers (framed boxes). Lines connecting *C. burkhardae* proteins to enzymatic functions are color-coded according to their E-values against the EggNOG database. The expression of three key genes (bold, orange) was confirmed with reverse-transcription PCR. (B) Expression of genes *KAA0152884*, *KAA0165906* and *KAA0166696* in *C. burkhardae*. Shown are the RT-PCR products obtained from three independent *C. burkhardae* cultures (labeled A-C), including the constitutively expressed aspartyl-tRNA synthetase gene (*AspRS*) as a positive control. Lanes 4, 8, 12 and 16 correspond to no-template controls.

An essential step in methionine biosynthesis is homoserine activation, which can be carried out by transfer of a phosphate group (Bartlem *et al.*, 2000), a *O-*succinyl group (Born and Blanchard, 1999), or an *O-*acetyl group (Goudarzi and Born, 2006) to the γ-hydroxyl group of homoserine. Our computational analysis did not identify homologs of a homoserine kinase or a homoserine *O-*succinyltransferase. Instead, we found two putative homologs of a homoserine *O-*acetyltransferase suggesting that, in *C. burkhardae,* homoserine is activated through the addition of an *O-*acetyl group (Fig. 1A and Table S1). The activated *O-*acetyl-homoserine is then converted to homocysteine, either directly or with cystathionine as an intermediate (Fig. 1A). However, the high sequence similarities between cystathionine-γ-synthase, cystathionine-β-lyase and cysteine-synthase made it difficult to predict which of the five candidate genes (KAA0166696, KAA0162384, KAA0146633, KAA0161056, KAA0169545) encoded which enzymatic activity, as previously discussed for cystathionine synthases and lyases in *T. pseudonana* (Bromke and Hesse, 2015). Nevertheless, genome analysis strongly suggests that *C. burkhardae* encodes a complete methionine biosynthesis pathway.

To test if the identified genes were transcribed, we performed reverse-transcription PCR on three of them (KAA0152884, KAA0165906 and KAA0166696) (Fig. 1B). KAA0152884 and KAA0165906 are predicted to encode a homoserine acetyltransferase, the key enzyme for homoserine activation, which is absent in methionine auxotrophs (Finkelstein, 1990; Ferla and Patrick, 2014; Bromke and Hesse, 2015). The gene product of KAA0166696 is predicted to act downstream of homoserine activation in methionine biosynthesis. As a control, we used the aspartyl tRNA synthetase (AspRS) gene whose expression has been demonstrated previously (Fischer and Hackl, 2016). We found that all tested genes were expressed in actively growing *C. burkhardae* cells (Fig. 1B). Based on the presence and expression of all essential methionine biosynthesis genes, we conclude that *C. burkhardae* is a methionine prototrophic organism.

### Density-based removal of food bacteria from cultures of *C. burkhardae*

In order to reduce potential background signal in our assays from the mixed bacterial community that is present as a food source in *C. burkhardae* cultures, we used a 10%/20% (w/v) iodixanol density cushion to separate bacteria from flagellate cells. Bacteria will penetrate the 20% iodixanol layer and pellet during centrifugation, whereas *C. burkhardae* cells collect at the 10%-20% iodixanol interphase. To evaluate the effectiveness of this process, cells from an untreated *Cafeteria* culture were compared to cells extracted from the 10%-20% iodixanol interphase. Cells were fixed and immobilized on Poly-L-Lysine-treated coverslips (PLL-treated), stained with 4′,6-diamidino-2-phenylindole (DAPI), and imaged with a fluorescence microscope. In untreated cultures, the bacterial DAPI fluorescence impaired the visualization of *C. burkhardae* DNA (Fig. S1A). When the culture was purified via iodixanol density cushion centrifugation, the bacterial concentration decreased sufficiently allowing the identification of individual cells of *C. burkhardae* (Fig. S1A). However, not all bacteria were removed during the procedure, probably owing to the heterogeneous species composition of the bacterial population, with some bacteria exhibiting buoyant densities similar to that of *C. burkhardae*.

We used flow cytometry to quantify the remaining bacteria in *C. burkhardae* cultures before and after density cushion centrifugation. As shown in Fig. S1B, bacterial populations can be distinguished from *C. burkhardae* cells in scatter plots. Gating and quantification (Table S2) of bacterial populations indicated that 20%-50% of bacteria from the original culture are still retained after purification, hence iodixanol density cushion centrifugation removed 50-80% of the bacteria (Fig. S1B) without impacting *C. burkhardae* populations. Thus, the procedure is adequate for reducing the bacterial background. Moreover, the additional centrifugation steps during cell preparation for fluorescent microscopy will further remove bacteria leading to levels that allow clear imaging of *C. burkhardae* cells.

### AHA is efficiently incorporated into*C. burkhardae* proteins

To test for AHA uptake in *C. burkhardae*, we incubated cultures that had been purified by density cushion centrifugation with different AHA concentrations for four hours. AHA incorporation in *C. burkhardae* proteins was initially analyzed by polyacrylamide gel electrophoresis (PAGE). Given the lack of protocols for protein extraction from *C. burkhardae,* we first tested chemical lysis conditions with different concentrations of the following nonionic detergents: Igepal^®^, Triton™ X-100, and NP-40. The best results were obtained with 1% Igepal^®^ (data not shown). Thus, after AHA incubation, *C. burkhardae* cells were incubated in a buffer containing 1% Igepal^®^ and the total and soluble protein fractions were loaded on a 12% SDS-PA gel to assess cell lysis. No differences were found between Coomassie-stained total and soluble fractions, indicating efficient lysis (Fig. S2). Next, soluble AHA-containing proteins were labeled by Click-iT^®^ with tetramethylrhodamine alkyne (TAMRA) dye and analyzed via SDS-PAGE. Coomassie staining confirmed that the same amount of protein was loaded in all the lanes, consequently, no differences are observed between samples (Fig. 2A, lanes 1-5). TAMRA detection showed that *C. burkhardae* cell lysates contained AHA-labeled proteins, when AHA concentrations of 0.25 mM to 2 mM were used (Fig. 2A lanes 2’-5’). TAMRA intensity was only slightly higher for 1 mM of AHA (Fig. 2A lane 4’) compared to the other AHA concentrations tested (45528 arbitrary unit [AU] counts for 0.25 mM AHA, 45600 AU counts for 0.5 mM AHA, 54344 AU counts for 1 mM AHA, and 48096 AU counts for 2 mM AHA) which could be caused by small differences during gel loading. As expected, the strongest Coomassie bands did not match the strongest TAMRA bands. This is due to Coomassie intensity being proportional to the amount of loaded protein while AHA incorporation depends on the number of methionine residues per protein.

**Figure 2.**
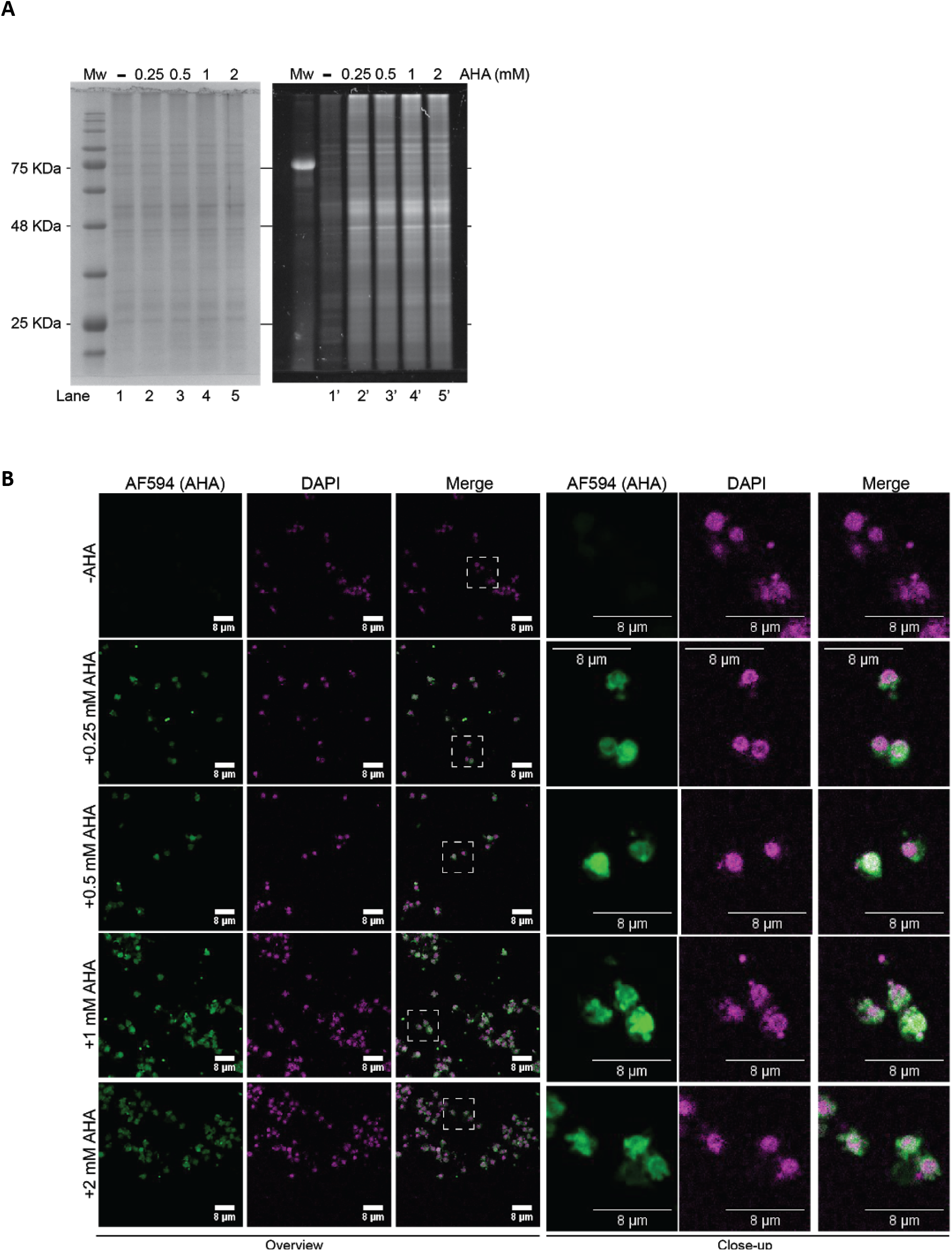
Incorporation of AHA in *C. burkhardae* proteins. (A) SDS-PAGE of soluble proteins extracted from 6×10^6^ *C. burkhardae* cells that were incubated with different concentrations of AHA. Gels were first stained with Coomassie (lanes 1-5) and then tested for the presence of AHA via coupling and excitation of a TAMRA fluorophore (lanes 1’-5’). (B) Confocal microscopy of *C. burkhardae* cells incubated with different concentrations of AHA. AHA residues were coupled to Alexa Fluor 594 and DNA was stained with DAPI. Areas outlined by white boxes are shown at higher magnification in the panels on the right.

In order to confirm that the AHA signal colocalized with *C. burkhardae* instead of residual bacteria, we labeled AHA residues with Alexa Fluor 594 (AF594) alkyne dye and analyzed cells using confocal microscopy. With all AHA concentrations tested, 100% of *C. burkhardae* cells showed a fluorescent signal in the AF594 channel (Fig. 2B), which indicates that AHA can replace methionine during protein synthesis in this protist. Most of the AHA-linked fluorescence signal was located around the nucleus and the mitochondria, which corresponds to the subcellular localization of protein biosynthesis. The highest signal was again obtained with 1 mM of AHA (fluorescence intensity 161 AU counts vs 126 AU counts for 0.25 mM AHA, 105 AU counts for 0.5 mM AHA and 123 AU counts for 2 mM AHA). However, the differences between AHA concentrations were small so that we decided to perform subsequent experiments with 0.5 mM of AHA in order to minimize potential stress for the cells.

### AHA is preferably incorporated into viral proteins during CroV infection

Next, we tested AHA incorporation into viral proteins during a *C. burkhardae* infection with the giant DNA virus CroV. We infected a *C. burkhardae* culture with CroV at a multiplicity of infection (MOI) of 2 and, after 1 h of incubation, removed the bulk of bacteria and free CroV particles by centrifugation on a 10-20% (w/v) iodixanol density cushion. As a control, we used a non-infected culture treated identically. After density cushion centrifugation, flagellate cells were resuspended in f/2 medium with 0.5 mM AHA or dimethyl sulfoxide (DMSO) (AHA-negative control) starting at 3 or 7 hpi. After 4 h of incubation with AHA, cells were harvested and lysed at 7 and 11 hpi, respectively. We analyzed the resulting protein extracts on a 12% SDS-PAGE stained with Coomassie Brilliant Blue R-250 (Fig. S3). The banding patterns were nearly identical between soluble and total fractions, indicating efficient cell lysis. Interestingly, only one Coomassie band was detected in the CroV-infected sample after 7 hpi that was not present in the uninfected sample (Fig. S3, arrow). An additional band appeared at 11 hpi (Fig. S3, asterisk). These two bands were identified by mass spectrometry as the major capsid protein CroV342 and the major core protein CroV332, respectively (Table S3). They are the two most abundant structural proteins in the CroV virion (Fischer et al, 2014) and good indicators for a productive CroV infection. Additional differences between infected and uninfected samples, as well as between 7 hpi and 11 hpi samples, were observed when we visualized AHA-containing proteins via TAMRA fluorescence (Fig. 3A, lanes 1’-4’ and 7’-10’). To compare these banding patterns directly with the CroV virion proteome, a protein lysate extracted from purified CroV particles is shown in lanes 5 and 6 of Fig. 3A. Several of these proteins were identified by mass spectrometry as CroV proteins (Figure 3A bands A to L, Table S3). Most of the proteins synthesized between 3 hpi and 7 hpi are probably non-structural CroV proteins because the banding pattern at 7 hpi differs from both the uninfected condition as well as from the purified CroV particle proteins (Fig. 3A lane 4’ vs lanes 2’ and 5). At 11 hpi, the TAMRA banding pattern resembles the Coomassie pattern of the CroV virion proteome (Fig. 3A lanes 10’ and 6), suggesting that most of the proteins synthesized in infected cells between 7 and 11 hpi are CroV virion components.

**Figure 3.**
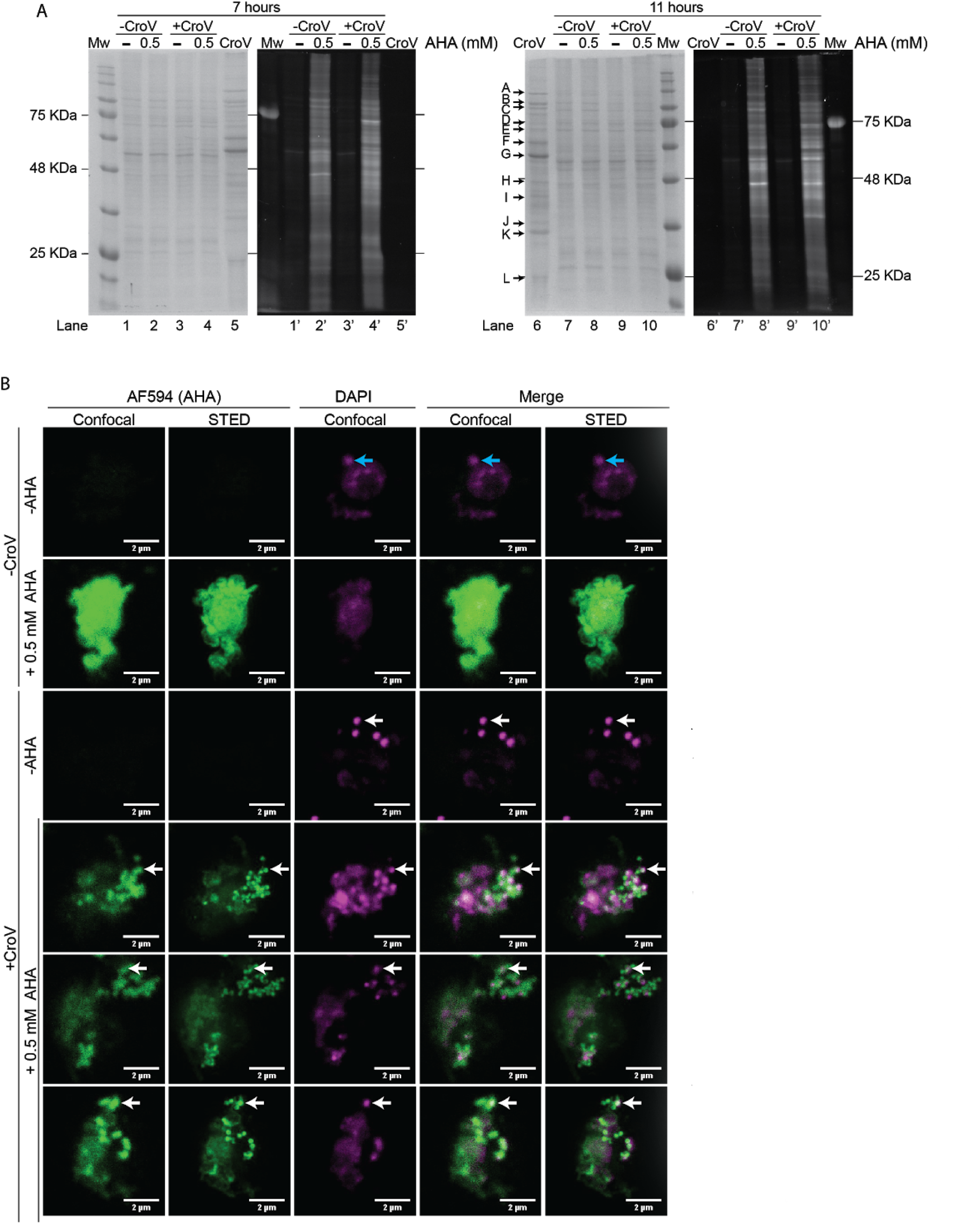
AHA incorporation during CroV infection. (A) SDS-PAGE of soluble proteins extracted from 6×10^6^ *C. burkhardae* cells that were either uninfected or infected with CroV, and incubated with or without AHA for four hours prior to harvesting at 7 hpi or 11 hpi. Gels were first stained with Coomassie (lanes 1-10) and then tested for the presence of AHA via coupling and excitation of a TAMRA fluorophore (lanes 1’-10’). Proteins extracted from purified CroV particles are shown in lanes 5 and 6, and bands that were analyzed by mass spectrometry are indicated by arrows A-L. (B) Confocal and STED microscopy of CroV-infected and non-infected *C. burkhardae* cells incubated with or without AHA. Cells were harvested at 7 hpi, AHA-containing proteins visualized by AF594 coupling and DNA stained with DAPI. White arrows indicate AF594 and DAPI signals of putative CroV particles, blue arrows depict putative mitochondria.

We then coupled the fluorescent dye AF594 to AHA moieties using Click-iT^®^ and analyzed uninfected and CroV-infected cells (7 hpi) by confocal microscopy. In uninfected cells, the AF594 signal was located around the nuclear and mitochondrial DAPI signal (DNA) (Fig. 3B). However, when *C. burkhardae* cells were infected with CroV, the AF594 signal overlapped largely with a brighter DAPI signal that drowned out the weaker nuclear and mitochondrial signals (Fig. 3B). The most intense DAPI spots had a diameter of ≈300 nm, which equals the diameter of a CroV capsid (Xiao *et al.*, 2017). In contrast to the well-defined DAPI structures, the AF594 signal obtained by confocal microscopy was rather diffuse (Fig. 3B). We, therefore, analyzed the same samples with Stimulated Emission Depletion (STED) microscopy, which resolved AF594 fluorescence into distinct dots that overlapped with the DAPI signal and had a diameter of 200-300 nm, strongly suggesting that they corresponded to CroV particles (Fig. 3B). Whereas all distinct DAPI dots had a matching AF594 signal, some AF594 dots were DAPI-negative. This indicates that the highest concentration of AHA-containing proteins is present in newly synthesized CroV particles, but not all particles present at 7 hpi contained DNA.

Our data show that AHA is efficiently incorporated into viral proteins during CroV infection of *C. burkhardae*. Assuming similar AHA acquisition rates of viral and eukaryotic proteins, this suggests an efficient metabolic rewiring of the infected cell towards viral protein production. The resulting virocell (Forterre, 2012) thus differs drastically in its proteome from uninfected cells (Fig. 3A). BONCAT labeling with AHA exploits this hallmark property of viral infection by preferentially labeling viral proteins during a specific phase of infection and in combination with STED microscopy, we show that visualization of individual CroV particles is possible. Although BONCAT has been applied to viruses such adenovirus (Banerjee *et al.*, 2011), vaccinia virus (Huang *et al.*, 2017) and Emiliania huxleyi virus (EhV207) (Pasulka *et al.*, 2018), we image individual giant virus particles for the first time using BONCAT in combination with super-resolution microscopy.

### AHA is incorporated in more than 95% of CroV particles

Next, we analyzed the percentage of AHA-positive CroV particles that were produced during the infection of *C. burkhardae* in the continuous presence of AHA. After virus-induced lysis, we removed cell debris by filtration and collected CroV particles on a 0.02 μm pore-size filter. AHA was labeled with Alexa Fluor 488 (AF488) alkyne dye by Click-iT^®^ reaction and the DNA was stained with DAPI. The samples were analyzed by non-confocal fluorescent microscopy. The AHA-negative control only showed a signal in the DAPI channel (Fig. S4A), whereas the AHA-treated sample contained AF488 positive viruses and bacteria. We manually quantified the AF488 and DAPI signals of approximately 2,000 CroV-like particles derived from three independent infection assays and found that 95%-100% of DAPI-stained CroV-like particles were also positive for AHA-AF488 (Table S4 and Fig. S4B). This indicates that AHA incorporation during CroV infection is highly efficient and reproducible. We also found that some CroV-like particles were not packed with DNA, as they were positive for AF488 fluorescence but negative in the DAPI channel (Table S4). The percentage of empty CroV-like particles varied from 1% in two experiments to 19% in the other experiment. This may indicate that CroV packaging efficiency varies between different infection experiments, or could reflect variance during microscopy quantification.

The highly efficient AHA labeling of CroV particles (>95%) is in concordance with results from other virus-host systems, for instance, an 87-94% labeling efficiency was reported for EhV207 (Pasulka *et al.*, 2018), >99% for herpes simplex virus (Serwa *et al.*, 2019) and 90% for vaccinia virus (Huang *et al.*, 2017). BONCAT can therefore be applied to a wide range of viruses.

### AHA incorporation does not affect CroV production or infectivity

To assess if the presence of AHA altered CroV replication, we analyzed by quantitative PCR (qPCR) the number of CroV genome copies produced during three independent CroV infections in the presence and absence of AHA. Two of the three biological replicates showed a 2.8-fold decrease in CroV DNA copy number in the presence of AHA compared to the AHA-negative control, whereas in the third replicate the viral DNA levels increased 1.5-fold in the AHA-treated sample (Fig. S5). These differences are not statistically significant, as determined by Wilcoxon signed-rank test analysis (p value = 0.75). Therefore, we conclude that an AHA concentration of 0.5 mM during CroV infection does not significantly affect CroV replication.

To test whether AHA-labeling might impair CroV entry in *C. burkhardae* cells, cultures were incubated with AHA-positive or AHA-negative CroV particles for 5 minutes and then fixed. AHA was labeled with AF594 alkyne dye by Click-iT^®^ reaction, DNA was stained with DAPI, and different lipid dyes for cytoplasmic membrane staining were tested, including Concanavalin A, CellBrite™ Fix and MemBrite™ Fix. Unfortunately, these dyes labeled inner membrane compartments of *C. burkhardae* both in live and fixed cells (data not shown). We therefore used an NHS ester linked to AF488 to label all proteins, allowing us to distinguish between extracellular and intracellular localization of CroV particles. As shown in Fig. 4A, infections with either native or AHA-containing CroV virions resulted in fluorescent signals of virus-like particles that were located inside *C. burkhardae* cells, indicating that the replacement of methionine by AHA does not affect CroV entry. Rapid fading of the AHA-linked AF594 signal in these samples, possibly caused by virion disassembly, prevented the use of high-resolution STED microscopy.

**Figure 4.**
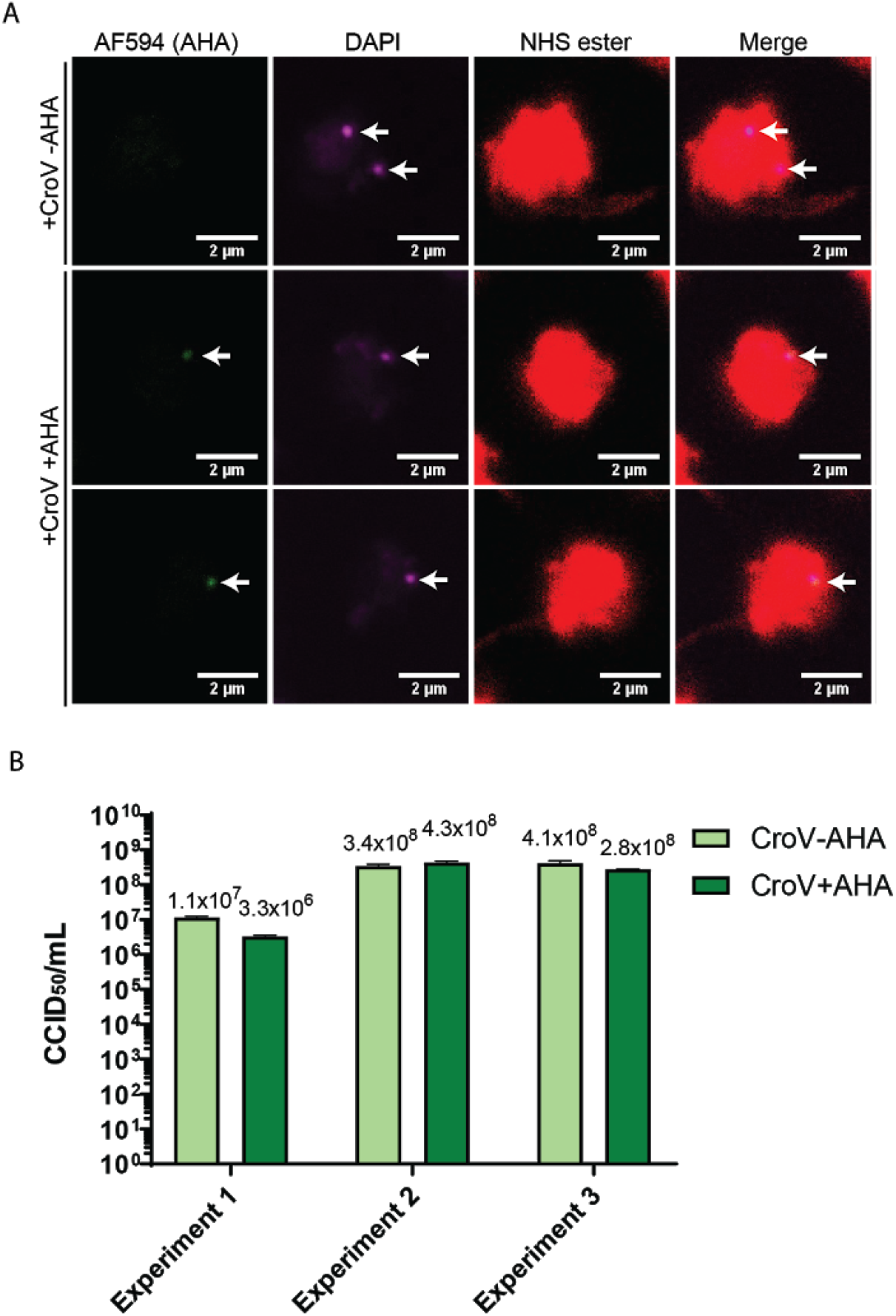
Infectivity assessment of AHA-labeled CroV particles. A) Confocal microscopy of early stages of CroV infection in *C. burkhardae* using AHA-labeled or unlabeled virions. *C. burkhardae* cultures were inoculated with native or AHA-labeled CroV particles at an MOI of 2 and incubated for five minutes prior to fixation. AHA residues were coupled to AF594, DNA was stained with DAPI, and flagellate cells were stained with AF488-NHS ester. Arrows point at AF594 and DAPI signals of virus-like particles. B) Infectivity comparison of native and AHA-labeled CroV particles. Infectivity values (CCID_50_/mL) of native (light green) or AHA-labeled (dark green) CroV particles derived from three independent infection experiments were determined by the end-point dilution method. Average values and SD (error bars) are shown for technical duplicates.

Finally, we compared the infectivity of AHA-labeled virions to native virions by end-point dilution assays using the same CroV lysates as for the qPCR test shown in Fig. S5. Based on genome copy quantification, the assay was started with same number of AHA-negative and AHA-positive CroV particles (1.7×10^6^ for experiment 1, 1.5×10^8^ in experiment 2 and 2×10^8^ in experiment 3). At three days post infection, we determined the infectious dose at which 50% of *C. burkhardae* cell cultures lysed (CCID50). The differences in CCID50 between AHA-negative and AHA-positive CroV particles were negligible in three independent assays (Fig. 4B), indicating that AHA has no apparent effect on CroV infectivity. This opens the possibility to study the CroV infection cycle with super-resolution microscopy using an antibody-free labeling method. In addition, this versatile, efficient, and non-toxic technique could be extended to the discovery and characterization of other giant viruses, even in environmental samples.

### Future perspectives

Although AHA and its alkyne analog L-homopropargylglycine (HPG) have been mostly used in methionine auxotrophic organisms to increase the efficiency of incorporation (Wisse *et al.*, 2017; Zhang *et al.*, 2017; Saleh *et al.*, 2019), its use has recently been extended to the analysis of natural bacterial communities, resulting in good incorporation in active methionine prototrophic bacteria (Hatzenpichler *et al.*, 2014; Leizeaga *et al.*, 2017; Couradeau *et al.*, 2019). In addition, four wild-type prototrophic eukaryotic organisms were reported to incorporate AHA or HPG: *Danio rerio* (Hinz *et al.*, 2012), *Caenorhabditis elegans* (Ullrich *et al.*, 2014), *Arabidiopsis thaliana* (Glenn *et al.*, 2017) and the coccolithophore *Emiliania huxleyi* (Pasulka *et al.*, 2018).

In this study, we demonstrated efficient incorporation of the methionine analog AHA in cellular and viral proteins during infection of the marine heterotrophic protist *C. burkhardae* with the giant virus CroV. We show that metabolic labeling can be used to visualize individual CroV particles, especially when imaged with super-resolution microscopy. Moreover, incorporation of AHA does not affect viral infectivity (Fig. 5), enabling downstream *in vivo* experiments with labeled particles. BONCAT labeling with AHA can thus be applied to study the infection cycles of giant viruses in a methionine prototrophic background.

**Figure 5.**
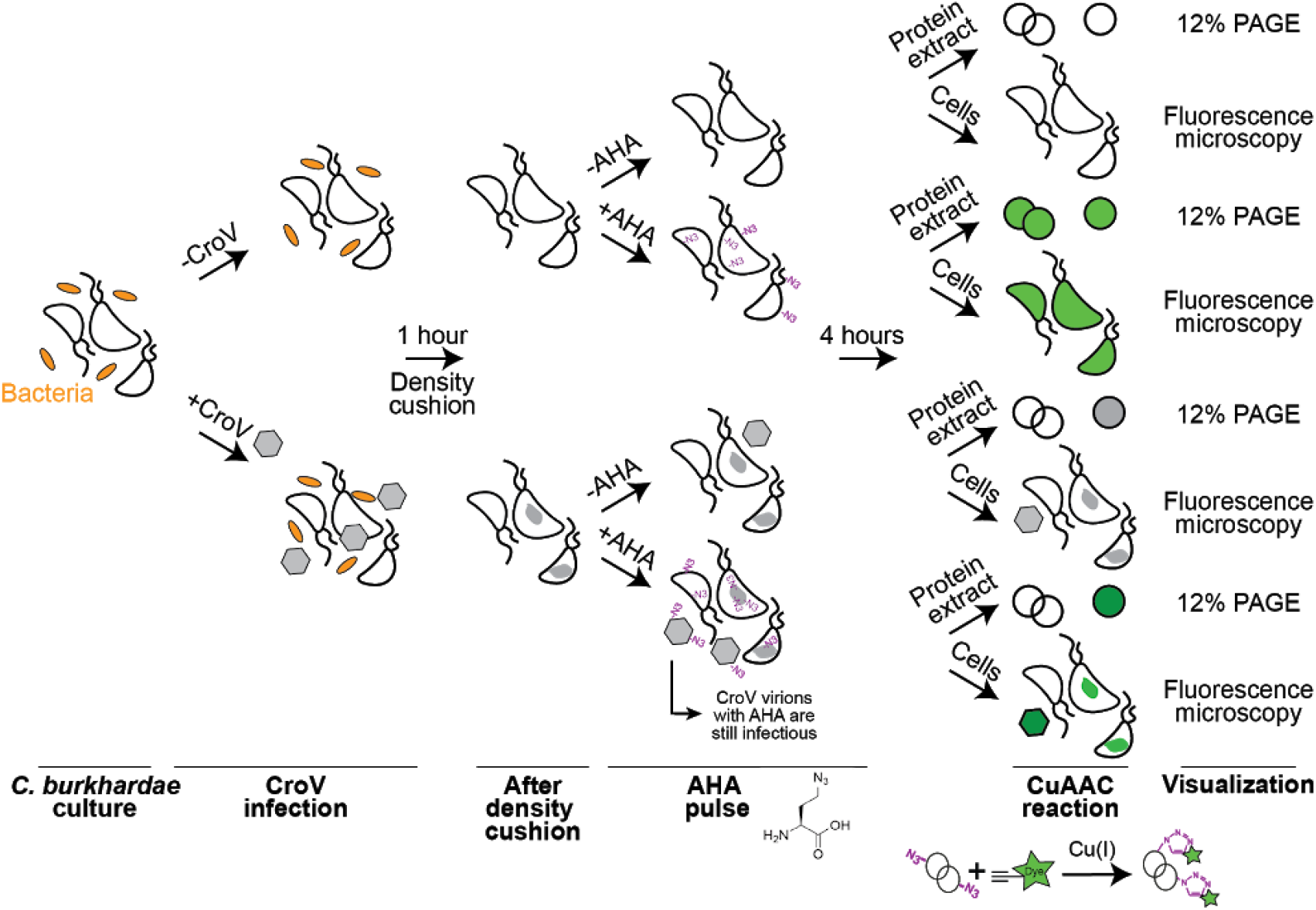
Experimental strategy for BONCAT labeling in *C. burkhardae*. Flagellate cultures were mock-infected or infected with CroV and incubated for one hour, followed by iodixanol density cushion centrifugation to remove bacteria and unattached CroV particles. Cells were then incubated for four hours with AHA or DMSO (negative control). AHA incorporation into nascent proteins was assessed by CuAAC (Click-IT)-mediated conjugation of AHA to fluorophores, either TAMRA alkyne dye for protein extracts and PAGE analysis, or AF594 or AF488 alkyne dyes for *in situ* cell analysis by fluorescence microscopy.

## Experimental procedures

### Host and virus strains

*Cafeteria burkhardae* (previously *C. roenbergensis*) RCC970-E3 strain was originally isolated from the South Pacific, approximately 2,200 km off the coast of Chile during the BIOSOPE cruise (Le Gall *et al.*, 2008). Suspension cultures were maintained as described before (Fischer and Hackl, 2016). Cultures for experiments were grown in flat-bottom 3 L polycarbonate Erlenmeyer flasks (VWR, Germany) at 22-25 °C. To determine the cell densities, 10 μL of a *C. burkhardae* culture were stained with 1 μL of Lugol’s acid iodine solution, loaded on a haemocytometer (Neubauer Chamber) and counted using a phase-contrast inverted microscope.

Cells were infected with CroV strain BV-PW1 (Garza and Suttle, 1995; Fischer *et al.*, 2010). For virus stock production, 500 mL of *C. burkhardae* RCC970-E3 were infected at an initial cell density of 5×10^5^ cells/mL with an MOI of 0.1. MOIs are based on CCID_50_/mL values calculated by the end-point dilution method described below. Cultures were incubated for 4 days until cells had lysed. Finally, cell debris was removed using a 1.2 μm filter.

### Iodixanol density cushion centrifugation

All centrifugation and incubation steps in this study were carried out at 20 °C unless specified otherwise. *C. burkhardae* RCC970-E3 cultures were centrifuged for 5 min at 4,500 g. Pellets were resuspended at a density of 2.5×10^8^ cells/mL in 1x PBS with 0.5 M NaCl. A total of 2.5×10^8^ cells were loaded in a SW40 ultra-clean centrifuge tubes (Beckman, Germany) containing 10% and 20% iodixanol density cushions (OptiPrep™, Sigma-Aldrich, Germany) cushion diluted in 1x PBS with 0.5 M NaCl, as described by (Duponchel *et al.*, 2019). Tubes were centrifuged for 20 min in a SW40Ti rotor at 20,000 rpm with slow braking in a Beckman Coulter Optima XE-90 ultracentrifuge. The interphase band containing *C. burkhardae* cells was recovered and washed with f/2 medium to remove residual iodixanol. After centrifugation for 5 min at 4,500 g, samples were resuspended in f/2 medium to a final concentration of 1×10^7^ cells/mL. Next, 5×10^6^ cells from treated and untreated conditions were centrifuged for 5 min at 4,500 g and the pellets were resuspended in 4% paraformaldehyde (PFA, Merck, Germany), and incubated for 20 min. After a 5 min centrifugation step at 7,000 g, pellets were washed with 1x PBS and centrifuged again. Pellets were resuspended in permeabilization buffer (3% (w/v) bovine serum albumin (BSA, Sigma-Aldrich, Germany) with 0.5% (v/v) Triton X-100 (Sigma-Aldrich, Germany) in 1x PBS) and incubated for 15 min. Cells were then immobilized by centrifuging for 5 min at 2,250 g onto poly-L-lysine (PLL)-treated coverslips, washed with 1x PBS and mounted on glass slides using ProLong ™ Diamond Antifade Mountant with DAPI (ThermoFisher, Germany). To obtain the PLL-treated coverslips, 15 mm-diameter coverslips were washed sequentially with 70% ethanol and H_2_O and treated with 0.01% (v/v) PLL solution (Sigma-Aldrich, Germany) for 20 min and drying overnight.

### Fluorescence microscopy

Imaging of bacterial backgrounds and quantification of CroV-AHA particles was performed on a Zeiss Axiovert 22M microscope with phase contrast and DAPI HC BP and GFP (for AF488) HC filter sets (F36-513 and F36-525 respectively, AHF Analysentechnik AG). For confocal and super-resolution microscopy, we used an Abberior easy 3D STED/RESOLFT QUAD scanning microscope (Abberior Instruments GmbH, Göttingen, Germany) built on a motorized inverted microscope IX83 (Olympus, Tokyo, Japan). The microscope was equipped with pulsed STED lasers at 595 nm and 775 nm, and with 355 nm, 405 nm, 485 nm, 561 nm, and 640 nm excitation lasers. Spectral detection was performed with avalanche photodiodes (APD) in the following spectral windows: 600-700 nm for AF594, 420-475 nm for DAPI, and 505-530 nm for AF488. Images were acquired with a 100x/1.40 UPlanSApo Oil immersion objective lens (Olympus). Pixel size was 200 nm for all overview pictures, 100 nm for close-up images in Fig. 2B, and 25 nm for images in Figs. 3B and 4A. For super-resolution imaging of AF594, the STED line at 775 nm was used. Image acquisition parameters were kept constant across the different samples to enable the quantification of signal intensities.

Microscopy images were processed and visualized with the ImSpector software package (Max-Planck Innovation) and/or ImageJ (imagej.nih.gov/ij/). Brightness and contrast settings were applied evenly to all images of an experiment.

### Flow cytometry

Seven independent cultures were analyzed, which were split into density cushion-purified or untreated batches with at least 10^7^ *C. burkhardae* cells each. The untreated batch was centrifuged for 10 min at 4,500 g and washed once in 1x PBS. After a second centrifugation step, the pellet was fixed with 4% PFA as described above. After a 5 min centrifugation at 7,000 g, cells were resuspended in 500 μl of 1x PBS and stored at 4 °C. The treated batch was centrifuged 10 min at 4,500 g and the pellet was subjected to iodixanol density cushion centrifugation as described above. The recovered cells were centrifuged for 10 min at 4,500 g and the pellet was incubated for 20 min at RT in 4% PFA. After centrifugation at 7,000 g for 5 min, treated samples were resuspended in 500 μl of 1x PBS and stored at 4 °C.

Cell count data were acquired on an LSR Fortessa X-20 cytometer (BD, Germany) and displayed in dot plots of SSC-A versus FSC-A on logarithmic axes (ranging from 10^−3^ to 10^5^). The FSC-A threshold was set to its minimum (200 r.u.) and a medium flow rate (approx. 35 μl/min) was used during acquisition. The sample injection port was cleaned between samples to prevent cross-contamination. Samples were filtered on cell-strainer cap (Falcon^®^, VWR, Germany), diluted 1:2 with 1x PBS and vortexed before acquisition. After stabilization of the flow rate, 10,000 total events per sample were acquired in technical duplicates.

Analysis of the raw data files was performed using FlowJo 10.5.3 software (BD, Germany). *C. burkhardae* cells, bacterial cells and background events were gated and the number of events for each cell population was obtained. The percentage of bacterial and flagellate cells corresponded to the average cell count in each treated sample divided by the average count of the respective untreated sample.

### Methionine synthesis genes in*C. burkhardae*

We used methionine synthesis proteins previously identified in the diatom *Thalassiosira pseudonana* (Bromke and Hesse, 2015) to identify candidate genes in *C. burkhardae* using BLAST 2.9.0+ (Altschul *et al.*, 1990). We then annotated those candidate genes against the EggNOG v4.5.1 (Huerta-Cepas *et al.*, 2016) database to confirm their putative functions (Table S1).

For extraction of total RNA from *C. burkhardae* RCC970-E3 culture, 500 μl of an exponentially growing suspension culture, which was previously submitted to iodixanol density cushion, were centrifuged for 5 min at 4,500 g, 4 °C. The supernatants were discarded and the cell pellets were immediately flash-frozen in N_2_ (l) and stored at −80 °C until further use. RNA extraction was performed with the Qiagen RNeasy Mini Kit (Qiagen, Hilden, Germany) following the protocol for purification of total RNA from animal cells using spin technology. Cells were disrupted with QIAshredder homogenizer spin columns (Qiagen, Hilden, Germany) and an on-column DNase I digest was performed with the Qiagen RNase-Free DNase Set (Qiagen, Hilden, Germany). RNA was eluted in 50 μl of RNase-free molecular biology grade water and DNA was removed by incubation with 1 μl TURBO DNase (2 U/μl) for 30 min at 37 °C according to the manufacturer’s instructions (Ambion via ThermoFisher Scientific, Germany). RNA samples were analyzed for quantity and integrity with a Qubit 4 Fluorometer (Invitrogen via ThermoFisher Scientific, Germany) using the RNA Broad Range and RNA Integrity and Quality kits, respectively.

For cDNA synthesis, 6 μl of each RNA sample were reverse transcribed using the Qiagen QuantiTect Reverse Transcription Kit according to the manufacturer’s instructions (Qiagen, Hilden, Germany). This protocol included an additional DNase treatment step and the reverse transcription reaction used a mix of random hexamers and oligo(dT) primers. Control reactions to test for gDNA contamination were performed for all samples by adding double-distilled (dd) H_2_O instead of reverse transcriptase to the reaction mix. The cDNA was diluted fivefold with RNase-free H_2_O and analyzed by PCR with gene-specific primers (Table S5).

PCR amplification was performed using 2 μl of the diluted cDNA in a 25 μl reaction mix containing 0.02 U of Q5^®^ High-Fidelity DNA Polymerase (NEB, Germany), 200 μM of dNTP, 5 μl of 5x Q5 High GC Enhancer and 0.5 μM of each primer in 1x of the Q5 Reaction Buffer. The PCRs were performed in a ProFlex PCR System (Applied Biosystems via ThermoFisher Scientific, Germany) with the following cycling conditions: 30 s denaturation at 98 °C, 35 cycles of 10 s denaturation at 98 °C, 20 s annealing at the temperature given by NEB TM calculator (https://tmcalculator.neb.com), and 25 s extension at 72 °C. The cycles were followed by a final extension period of 2 min at 72 °C. For product analysis, 1 μl of each reaction was loaded on a 1% (w/v) agarose gel supplemented with GelRed. The marker lanes contained 0.5 μg of GeneRuler 1 kb DNA ladder (Fermentas, ThermoFisher Scientific, Germany). The gel was run for 1 h at 100 V and visualized on a ChemiDoc MP Imaging System (BioRad, Germany).

### AHA labeling of *C. burkhardae* and CroV

To study AHA incorporation in newly synthesized *C. burkhardae* proteins, different AHA concentrations were tested. First, 2.5×10^8^ cells were purified by iodixanol density cushion centrifugation as described above. Recovered cells were resuspended in f/2 medium (without yeast extract) at a density of 10^7^ cells/mL. For each sample, 1.5×10^7^ cells were incubated for 4 hours with 3 μL of AHA diluted in DMSO (Acros Organics via ThermoFisher Scientific, Germany) to final concentrations of 0, 0.25, 0.5, 1 or 2 mM AHA. After incubation, samples were prepared for SDS-PAGE and microscopy analysis as described below.

To analyze AHA incorporation in CroV proteins, 1 L of *C. burkhardae* RCC970-E3 suspension culture was diluted to 4×10^5^ cells/mL on the day before the experiment. A total of 2.5×10^8^ cells were then infected with CroV at an MOI of 2 (stock quantified by end-point dilution). The mock infection received f/2 medium instead of CroV. After a 1 hour incubation, bacteria and free CroV particles were removed by density cushion centrifugation. Recovered cells were resuspended in f/2 medium at a density of 1×10^7^ cells/mL and incubated for a total infection time of 7 or 11 hours. At 3 and 7 hours post infection, 0.5 μL of 1 M AHA (0.5 mM final concentration) and 0.5 μL of DMSO (negative control) were added, respectively, resulting in a 4 hour incubation time prior to harvesting.

AHA incorporation was assessed by SDS-PAGE using 6×10^6^ cells. Cells were centrifuged for 6 min at 5,000 g after AHA incubation. Pellets were washed with 500 μL of 1x PBS and centrifuged again. Samples were frozen at −80 °C until use. To extract *C. burkhardae* and CroV proteins, pellets were resuspended in 50 μL of lysis buffer (20 mM Tris-HCl pH 7.5, 1 mM EDTA, 0.5 mM Na3VO4, 1 mM benzamidine hydrochloride (Sigma-Aldrich, Germany), 5x *cOmplete*™ EDTA-free protease inhibitor cocktail (Roche, Germany), 10% (v/v) glycerol, 1% (v/v) Igepal^®^ (Sigma-Aldrich, Germany) and 100 μM phenylmethylsulfonyl fluoride (Sigma-Aldrich, Germany) in H_2_O) and incubated 20 min on ice. Lysis was checked by phase-contrast inverted microscopy and by comparing the SDS-PAGE banding pattern of 5 μL of the lysate (total protein fraction) and 5 μL of the lysate after centrifuging for 15 min at 12,000 g, 4 °C (soluble protein fraction). After cell lysis, incorporated AHA was detected by labeling 40 μL of each soluble protein fraction with 22.5 μM of TAMRA alkyne in a final volume of 128 μL following the Click-IT^®^ Protein Reaction Buffer Kit recommendations (ThermoFisher, Germany). After methanol/chloroform protein precipitation, protein pellets were resuspended in 20 μL of 1x Laemmli buffer (10 min shaking followed by 10 min incubation at 70 °C) and 2 μL were loaded on a 12% SDS-PA gel. After electrophoresis at 150 V for 1.5 hours, gels were stained with InstantBlue^®^ Coomassie (Expedeon via Abcam, Germany) and visualized on a ChemiDoc MP Imaging System using rhodamine settings for TAMRA detection and Coomassie settings for Coomassie detection.

For microscopy, 3×10^6^ cells were centrifuged for 6 min at 5,000 g and fixed and permeabilized as described above. After pelleting the cells for 5 min at 7,000 g, AHA was labelled with 0.5 μM AF594 alkyne dye following the Click-iT^®^ Cell Reaction Buffer Kit instructions. Cells were centrifuged for 5 min at 7,000 g and washed twice with 200 μL of 1x PBS containing 3% BSA and 0.1% (v/v) Tween-20 (BioRad, Germany). Mounting of cells for STED microscopy was performed as described above.

### Production and analysis of AHA-labeled CroV

Three independent batches of AHA-containing and AHA-free CroV particles were produced and further analyzed by fluorescence microscopy, qPCR and infectivity assays. For each batch, 2×10^7^ CroV-infected *C. burkhardae* cells at a density of 1×10^7^ cells/mL were density cushion-purified and incubated for 24 hours with 1 μL of 1 M AHA (0.5 mM final concentration) or 1 μL of DMSO for the negative control. After cell lysis, samples were 1.2 μm filtered and 100 μL of the filtrate were fixed with 32.5 μL of 16% PFA (4% final) for 20 min. Samples were immobilized on a 25 mm Anodisc Whatman filter with 0.02 μm pore size, as described previously (Berjón-Otero *et al.*, 2019) and permeabilized for 15 min. AHA residues were coupled to 0.5 μM AF488 alkyne dye using the Click-iT^®^ Cell Reaction Buffer Kit. Filters were washed three times with 1 mL of 3% BSA with 0.1% Tween-20 in 1x PBS and mounted on slides as described above. CroV particles positive for DAPI and AF488 were manually counted using ImageJ software.

For the CroV cell entry assay, 5×10^6^ density cushion-purified *C. burkhardae* cells were resuspended with AHA-containing or AHA-free CroV particles at MOI 2 (based on qPCR). After 5 min of incubation, cells were fixed, permeabilized, and immobilized as described before. AHA was labeled with 0.5 μM AF594 alkyne dye using the Click-iT^®^ Cell Reaction Buffer Kit. Coverslips were washed three times with 200 μL of 3% BSA with 0.1% Tween-20 in 1x PBS and once with 1x PBS. *C. burkhardae* cells were stained 2 min at RT with 40 μg/mL AF488 NHS ester (ThermoFisher, Germany). Prior to mounting and STED microscopy, coverslips were washed three times with 1x PBS.

### Quantitative PCR

Viral genomic DNA (gDNA) was extracted from 100 μl of viral suspension complemented with 100 μl of 1x PBS with the QIAamp DNA Mini kit (Qiagen, Hilden, Germany) following the manufacturer’s instructions for DNA purification of total DNA from cultured cells, with a single elution step in 100 μl of ddH_2_O and storage at −20 °C. 1 μl of gDNA was used as template in a 20 μl qPCR reaction containing 10 μl of 2x Fast-Plus EvaGreen Master Mix with low ROX dye (Biotium, Inc. via VWR, Germany), 10 pmol of each forward and reverse primer (Table 1), and 8.8 μl of ddH_2_O. No-template controls (NTC) contained ddH_2_O instead of gDNA. Each qPCR reaction (sample, NTC, or standard) was performed in technical duplicates, with individual replicates differing in their quantification cycles (Cq) by about 0.08% on average (0.084% ± 0.086%, n = 6). The limit of detection for this assay was ≈10 copies, which equates to about 5,000 copies/mL of suspension culture. The Cq values of the NTC controls were consistently below the limit of detection. Thermal cycling was performed in a Stratagene Mx3005P qPCR system (Agilent Technologies, Germany) with the following settings: 95 °C for 5 min, 40 cycles of 95 °C for 10 s followed by 60 °C for 25 s and 72 °C for 25 s, a single cycle of 72 °C for 5 min, and a final dissociation curve was recorded from 50 °C to 95 °C. qPCR results were analyzed using MxPro qPCR software version 4.10 (Stratagene via Agilent Technologies, Germany). The threshold fluorescence was set using the amplification-based option of MxPro software. Standard curves were calculated from a tenfold dilution series that ranged from 10^1^ to 10^8^ molecules of gDNA extracted from a known amount of CroV particles, the concentration of which had been determined by epifluorescence microscopy. The target sequence crov283 gene (GenBank accession number ADO67316.1) was amplified by primers CroV-qPCR-9 and CroV-qPCR-10 (Table S5). Using these primers, the R2 value for the standard curve was 0.982, the amplification efficiency was 115.0%, and the standard curve equation was y = −2.995log(x) + 40.91.

### CroV infectivity assays

End-point dilution assays for measuring infective titers of CroV have been described previously (Fischer and Hackl, 2016). Plates were inspected for cell lysis at 5 days post infection by phase-contrast microscopy. For the infectivity comparison of AHA-labeled *versus* AHA-free CroV virions, dilutions ranged from 10^−3^ to 10^−9^ and plates were analyzed at 3 days post infection. The statistical method by Reed and Muench (Reed and Muench, 1938) was used to determine the CCID_50_/mL.

### Mass spectrometry

To identify SDS-PAGE protein bands, peptide map fingerprinting (PMF) was performed using standard procedures, including protein treatment with dithiothreitol and iodoacetamide followed by trypsin digestion as described by Wulfmeyer (Wulfmeyer *et al.*, 2012). Briefly, peptides eluted from gel slices were purified on ZipTip C18 columns (Millipore, Germany) and applied to a stainless steel target together with α-cyano-4-hydroxycinnamic acid as a matrix. The peptides were analyzed in reflectron mode using a Shimadzu Biotech Axima Performance MALDI-TOF mass spectrometer. Calibration was via nearest neighbor external standards, using 8 peptides (Sigma Aldrich) with m/z ratios from 757.4 to 3657.9. Mass lists from the individual PMF spectra were submitted to an in-house Mascot Server PMF search engine using the NCBI nr database, limiting the taxonomic range to *C. burkhardae* and CroV. Additional search parameters were set to monoisotopic mass, charge 1+, maximum of 1 missed cleavage, peptide tolerance of 0.3 m/z, and p<0.05. The root mean square (RMS) errors on the peptide mass matches ranged from 21–102 ppm. As a control, all searches were repeated using the decoy database generated by the Mascot Server software, using the same settings.

## Acknowledgments

This work was supported by the Max Planck Society, the Gordon & Betty Moore Foundation (grant #5734), and a European Molecular Biology Organization (EMBO) long-term fellowship to M.B.-O. (ALTF-726-2017). We are grateful to C. Suttle for access to host and virus strains, and to the Roscoff team for maintaining and distributing protist strains. We thank A. Weinmann for technical assistance, E. D’Este for support with confocal and STED microscopy, S. Fabritz, M. Müller and C. Ulrich for mass spectrometry, C. Roome for IT support and I. Schlichting for mentoring and support.

The authors declare no conflict of interest.

