## Supplemental Information for "Visualization of giant virus particles using BONCAT labeling and STED microscopy"

<u>Content:</u>

Supplemental Figures 1-5 with legends

Supplemental Table 5 with legend

Legends for Supplemental Tables 1-4

For Supplemental Tables 1-4, see accompanying Excel file

**Figure S1**

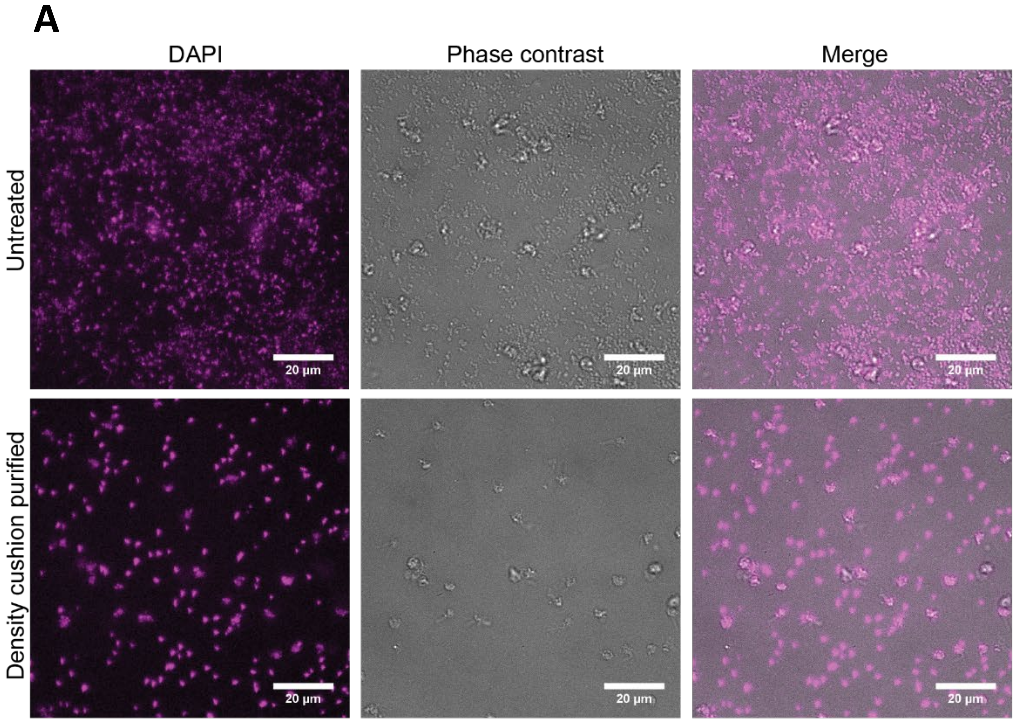


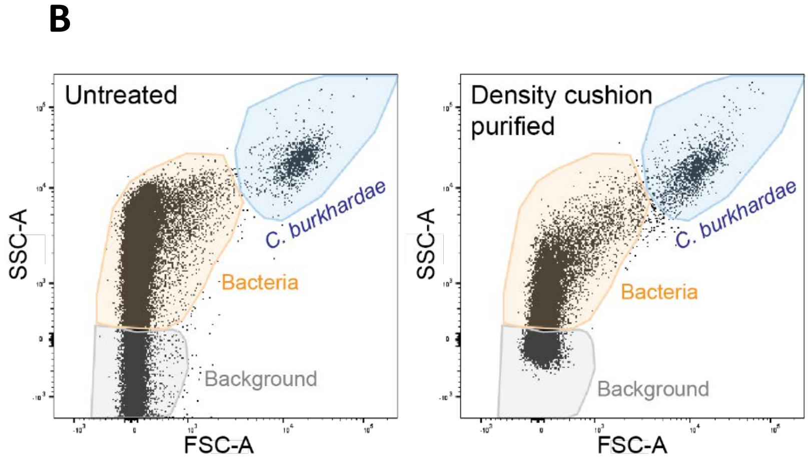


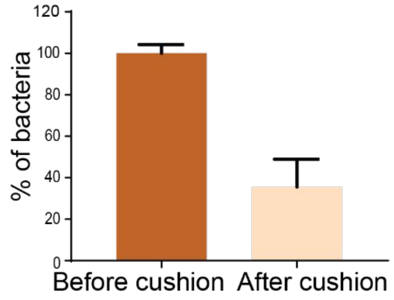


**Figure S1. Separation of *C. burkhardae* cells and bacteria by iodixanol density cushion centrifugation.** (A) Microscopy images of a *C. burkhardae* culture containing a mixed bacterial community before (top) and after (bottom) centrifugation onto a 20% (w/v) iodixanol cushion**.** DNA was stained with DAPI (first column). (B) Flow cytometry analysis of bacteria and flagellate populations in a mixed *C. burkhardae* culture before and after iodixanol density cushion centrifugation. The cytograms show side scatter and forward scatter of 10,000 events (top). The bottom graph shows relative bacterial abundances based on the quantification of events in the bacterial gate. See also Table S2.

### Figure S2

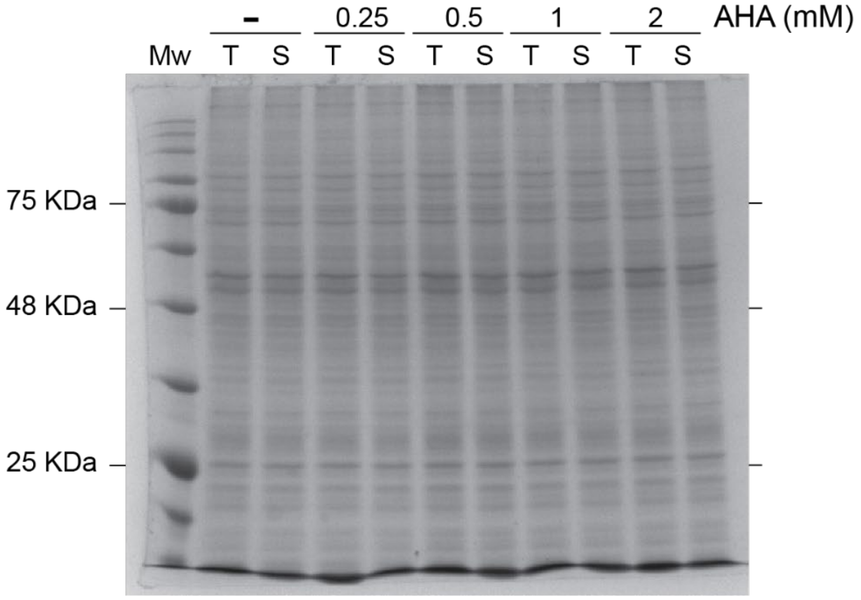


**Figure S2. SDS-PAGE of total and soluble protein fractions obtained from a lysate of *C. burkhardae* cultures incubated with different AHA concentrations.** $6x10^6$ cells of *C. burkhardae* culture, incubated with 0, 0.25, 0.5, 1 or 2 mM of AHA, were incubated in lysis buffer. Five microliters of the lysate supernatant (total protein fraction, T) and 5 µL of the lysate after centrifuging at 12,000 g, 15 min, 4 °C (soluble protein fraction, S) were resolved by 12% SDS-PAGE and Coomassie staining.

**Figure S3**

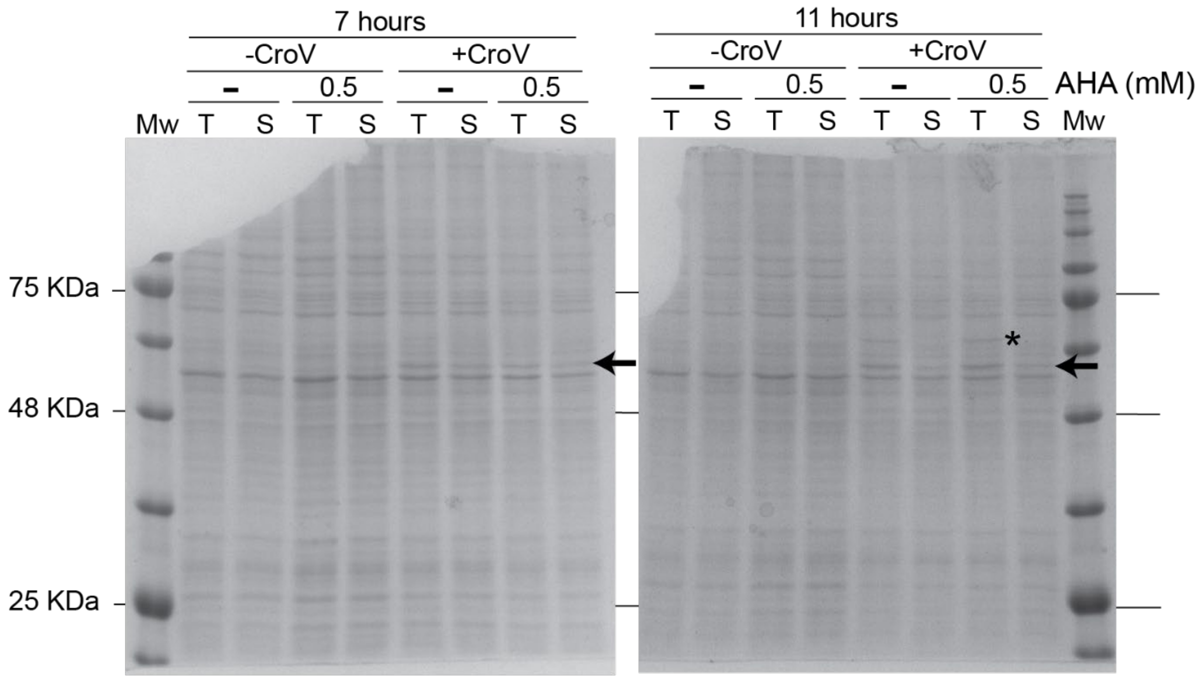


**Figure S3. SDS-PAGE of total and soluble protein fractions obtained from a lysate of CroV-infected and non-infected *C. burkhardae* cultures incubated in presence or absence of AHA.** 6x$10^6$ cells of CroV-infected (MOI 2) or non-infected *C. burkhardae* culture, incubated with 0 or 0.5 mM of AHA, were incubated in lysis buffer. 5 µL of the lysate supernatant (total protein fraction, T) and 5 µL of the lysate after centrifuging at 12,000 g, 15 min, 4 °C (soluble protein fraction, S) were resolved by 12% SDS-PAGE and Coomassie staining. The left gel shows the protein extract at 7 hpi, the right gel at 11 hpi. Arrows point at a CroV-specific band that was present at both timepoints, whereas the asterisk points at a band that was present at 11 hpi only. These bands were analyzed by mass spectrometry, see table S3.

### Figure S4

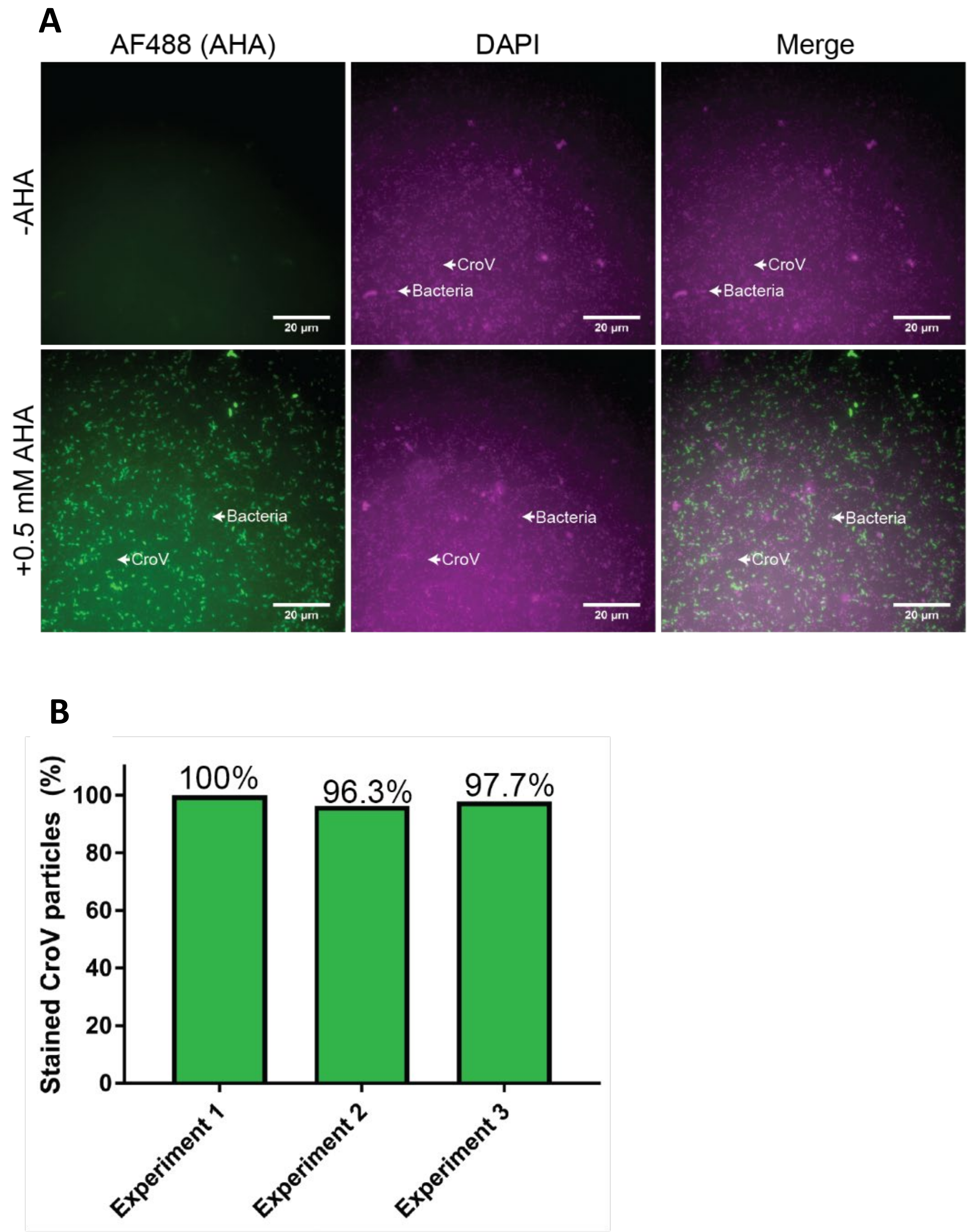


**Figure S4. Efficiency of CroV labeling with AHA.** A) Fluorescence microscopy of native and AHA-labeled CroV particles. Proteins containing AHA were visualized by coupling and exciting AF488 and DNA by DAPI staining. B) Percentage of AHA-positive CroV particles obtained from three independent labeling experiments. AHA was coupled to AF488, DNA was stained with DAPI (see Fig. S4A), and virus-like particles (VLPs) were counted in each channel by fluorescence microscopy. Shown is the percentage of DAPI-positive VLPs that also stained positive for AF488. See Table S4 for numerical data.

### Figure S5

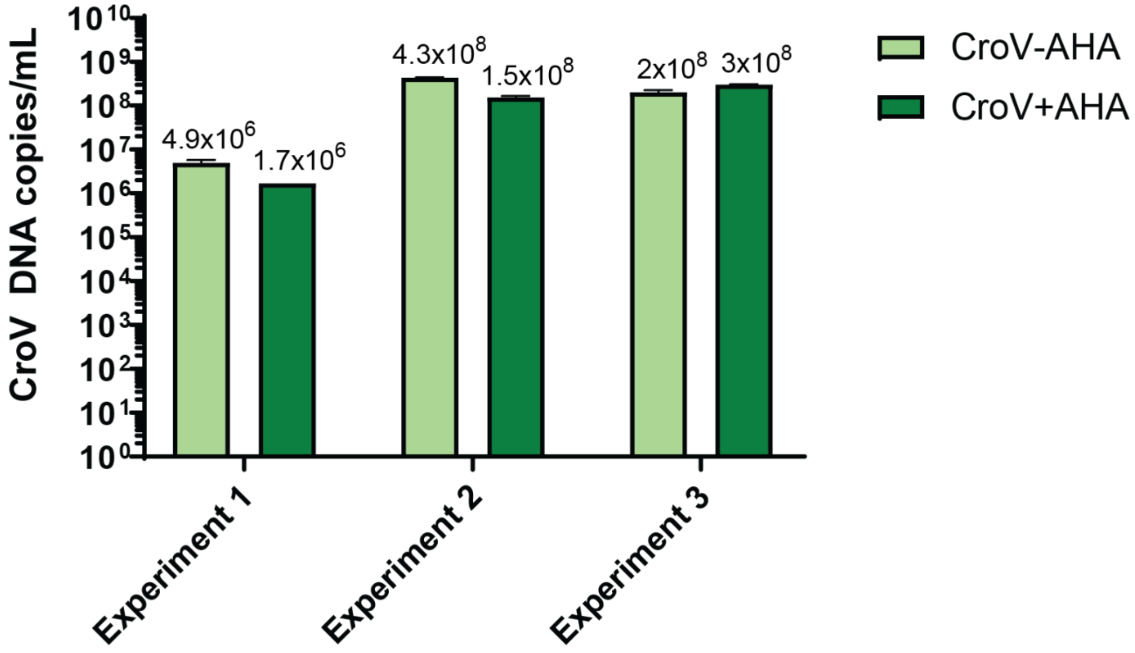


**Figure S5. CroV DNA copy numbers from three independent stocks of native CroV and CroV containing AHA.** DNA copy numbers were estimated by qPCR using primers CroV-qPCR-9 and CroV-qPCR-10 (Table S5).

**Table S5. Primers used in this study.**

| ***C. burkhardae* Gene ID** | **Primer name** | **Primer sequence (5'-3')** |
|---|---|---|
| KAA0152884 | HSATg1192-F | GAGGTGCGCGTGACTGTGGA |
| | HSATg1192-R2 | CCGCCCATGGCGAACTTGCT |
| KAA0165906 | HSATg6921-F | ACGTCGCAGGTACGCATTGG |
| | HSATg6921-R1 | TCGCTGTCCACGCCCACAAC |
| KAA0166696 | OACPTg1315-F | TCAAGGACACCAAGCATGCGG |
| | OACPTg1315-R1 | AGGAAGGCGGCGAGCTCTGT |

### Legends for Supplemental Tables 1-4 provided in Excel format

**Table S1. *C. burkhardae* genes predicted to be involved in the methionine pathway.**

**Table S2. Number of events detected in bacteria and *C. burkhardae* populations by flow cytometry, before and after iodixanol density cushion centrifugation.** This table shows the quantification of seven independent experiments. 'Replicate' refers to a technical duplicate where the same sample is analyzed again by flow cytometry. The percentage of bacteria corresponded to the average of bacteria number from each sample divided by the average of the corresponding 'Culture' sample. When the analysis was not performed in technical duplicate, the number of bacteria was taken directly for the calculation. The same calculation was done with the number of *C. burkhardae* cells to obtain the ratio of protists.

**Table S3. CroV proteins identified by mass spectrometry.**

**Table S4. Quantification of the CroV particles positive for AHA and/or DAPI signals in three independent experiments.** This table indicates the number of CroV particles that present AF488 (staining AHA) and DAPI signal from around 1,700-2,200 CroV particles. Counting was manually done using ImageJ software. The percentage of CroV particles containing AHA was estimated by comparison of the CroV particles positive for DAPI and for AF488. CroV particles with only AF488 signal were considered as DNA empty particles and were not taken into account in the percentage of AHA positive particles calculation. Percentage of empty CroV particles (AHA positive, DAPI negative) was calculated as relative to AHA positive CroV particles (100%).
